# The Evolution and Fate of Diversity Under Hard and Soft Selection

**DOI:** 10.1101/2020.05.13.091124

**Authors:** Patrick J. Chen, Rees Kassen

## Abstract

How genetic variation arises and persists over evolutionary time despite the depleting effects of natural selection remains a long-standing question. Here, we investigate the impacts of two extreme forms of population regulation – at the level of the total, mixed population (hard selection) and at the level of local, spatially distinct patches (soft selection) – on the emergence and fate of diversity under strong divergent selection. We find that while the form of population regulation has little effect on rates of diversification it can modulate the long-term fate of genetic variation, diversity being more readily maintained under soft selection compared to hard selection. The mechanism responsible for coexistence is negative frequency dependent selection which, while present initially under both forms of population regulation, persists over the long-term only under soft selection. Importantly, coexistence is robust to continued evolution of niche specialist types under soft selection but not hard selection. These results suggest that soft selection could be a general mechanism for the maintenance of ecological diversity over evolutionary time scales.

## Introduction

Ecologists and evolutionary biologists have long struggled to reconcile the abundant diversity among ecologically equivalent (i.e., those competing for the same resources) genotypes or species in nature with the expectation that natural selection should eliminate all but the fittest type. Indeed, we are still some way from a rigorous, empirically validated account of the evolution and maintenance of diversity, in part because most theory on diversity in population genetics and ecology concerns the conditions necessary to *maintain* it, not its *origin* (1).

The leading explanation is the ecological theory of diversification, which sees diversity as the result of strong divergent selection resulting in the evolution of locally adapted niche specialists (2,3). Commonly, divergent selection is thought to be generated by spatial variation in the conditions of growth such that different habitats or patches in the environment favour different optimal phenotypes. Provided dispersal among patches is weak relative to the strength of selection, niche specialists that have high fitness in some patches and low fitness in others evolve readily and can be maintained over time (2–9). Environmental variation in time, by contrast, generates fluctuating selection that favours a single, broadly adapted generalist type and so the conditions for the evolution and maintenance of diversity are far more restrictive (10).

This account of the generation and maintenance of diversity is appealing both for its generality, applying with equal force to haploids and diploids (or any other ploidy level), unicellular or multicellular organisms, and asexual or sexual modes of reproduction. It is not complete, however. Population genetic models for the maintenance of genetic polymorphism in spatially variable environments, which have been intensively studied for decades (8,10–12), identify two further conditions are required for the stable maintenance of diversity (i.e. – a protected polymorphism). The first is that niche specialist types exhibit a fitness trade-off across patches. In other words, the rank of relative fitness has to change across patches such that no single genotype is superior across all conditions of growth. The second is that population size must be regulated at the level of the local patch, before formation of a globally mixed dispersal pool. When this happens, each patch is guaranteed to contribute offspring to the next generation. Such local population regulation generates negative frequency-dependent selection that protects rare specialists from being lost (13; also called ‘soft selection’ by 14). If population regulation occurs at the level of the global dispersal pool (termed ‘hard selection’), more productive patches overwhelm dispersal from less productive patches, resulting in the eventual replacement of the type that is fittest in the niche that contributes most to the population (14–18; see also 19). Diversity can, therefore, be maintained under soft selection but not under hard selection, provided the fitness trade-offs across environments are fixed or evolve such that intermediate phenotypes have lower fitness than expected from a linear trade-off function (20–22).

Evidence bearing on the role of spatial variation in driving the emergence, coexistence, and long-term fate of diversity, as well as the robustness of trade-offs to continued selection, is mixed. While there is abundant experimental evidence that divergent selection leads to the evolution of specialization and fitness trade-offs in the absence of dispersal (reviewed in Kassen 2014) and a handful of studies showing that high rates of dispersal can prevent ecological diversification in spatially variable environments (23–25), far less attention has been paid to the conditions governing coexistence and the long-term fate of diversity. In fact, we are aware of just two direct tests of the role of soft and hard selection, both limited to ecological time scales. Bell (26) found that the manner of population regulation had little impact on the quantity of genetic variation in fitness maintained in a genetically diverse population of the unicellular algae, *Chlamydomonas reinhardtii*. Gallet et al (2018), on the other hand, found support for the predictions of the theory: reciprocally marked genotypes of *Escherichia coli* resistant for tetracycline and nalidixic acid, respectively, were maintained under soft but not hard selection when both drugs were delivered at sub-inhibitory concentrations in different patches (27). It remains unclear whether Gallet et al’s results are robust to prolonged selection on evolutionary time scales.

We have previously documented the emergence and coexistence of antibiotic resistant and sensitive genotypes from an initially isogenic population of *Pseudomonas aeruginosa* distributed into a two-patch environment connected by dispersal where one patch was supplemented with drug and the other was not (28). Despite the fact that population regulation was not directly manipulated in this experiment, resistant and sensitive types coexisted in the experiment due to a fitness trade-off between growth rate and resistance (sensitive strains grow faster than resistant ones in the absence of drug) that was underlain by negative frequency-dependent selection. Interestingly, the long-term fate of diversity was governed by the cost of resistance: compensatory mutations reducing the cost of resistance led to the gradual replacement of sensitive types by resistant ones, albeit at a rate far slower than that observed for temporal varying or constant antibiotic selection. This result suggests that negative frequency-dependent selection is not sufficient to support diversity indefinitely in the face of continued selection.

Here we use the same experimental set up to evaluate how the scale of population regulation impacts the evolution and fate of diversity in spatially variable environments. We track diversification in 12 independently evolved isogenic lines of the opportunistic pathogen *Pseudomonas aeruginosa* strain PA14 propagated in environments that mimic as closely as possible the two-patch model with dispersal outlined above under either hard or soft selection. Selection occurs due to the addition of 0.3 μg/ml of the fluoroquinolone antibiotic ciprofloxacin to standard Luria-Bertrani (LB) medium, a concentration sufficient to reduce population densities to 20% that of drug-free conditions. Controls consist of a permissive (PERM) environment where both patches contain drug-free Luria-Bertrani (LB) medium or a selective (SEL) environment where both patches contain LB supplemented with ciprofloxacin. Spatially variable environments are created, as in Leale and Kassen (2018), by adding ciprofloxacin to one but not the other patch and allowing dispersal between patches by mixing aliquot during serial transfer. The manner of population regulation is manipulated by adjusting the population density during dispersal: hard selection (HARD) is emulated by mixing equal volume aliquots from a pair of patches without regard to the relative density of cells in each subpopulation; soft selection (SOFT) at the level of the dispersal pool is imposed by ensuring the density of cells contributed by each patch is equal prior to dispersal. Our experiment is fully factorial, comprising 3 selection environments × 2 forms of population regulation × 12 replicate populations = 72 evolving populations, and propagated for 40 days, or ~ 264 bacterial generations.

## Results

Resistance, defined as a minimum inhibitory concentration (MIC; the lowest drug concentration that completely inhibits growth of the ancestral strain) of >2 g/ml ciprofloxacin, failed to evolve under the PERM treatment but evolved and spread rapidly under SEL conditions, as expected (**Supp. Fig. 1**). Population regulation had no detectable impact on the evolution and spread of resistance under either condition. We see strikingly different results under divergent selection, where one patch is supplemented with drug and the other is not, where resistance and sensitive types evolve rapidly and coexist at intermediately frequencies under both forms of population regulation (**Fig. 1**). Inspection of figure 1 reveals that there is little difference between treatments in the rate at which resistance evolves but the final average frequency of resistance was marginally higher under hard selection than soft selection, a result confirmed by fitting a two-parameter logistic growth model fit to the data that estimates the maximum rate of increase and final frequency of resistance (analogous to the intrinsic rate of growth and carrying capacity in population growth models; rate of increase: ***F***_***1,19***_ **= 0.197, *P* = 0.662**; final frequency ***F***_***1,19***_ **= *P* = 0.030**). Notably, the dynamics of resistance among independently evolved lines experiencing divergent selection are highly variable, a result we have previously shown to be due to high rates of clonal interference due to large population sizes (~2 × 10^8^ CFUs, or colony forming units, per ml) and so high mutation supply rates (28). This variation notwithstanding, our results suggest that population regulation has little effect on the rate of diversification but can modulate the long-term fate of coexisting types in a manner consistent, in direction at least, with the predictions of theory.

**Figure 1.**
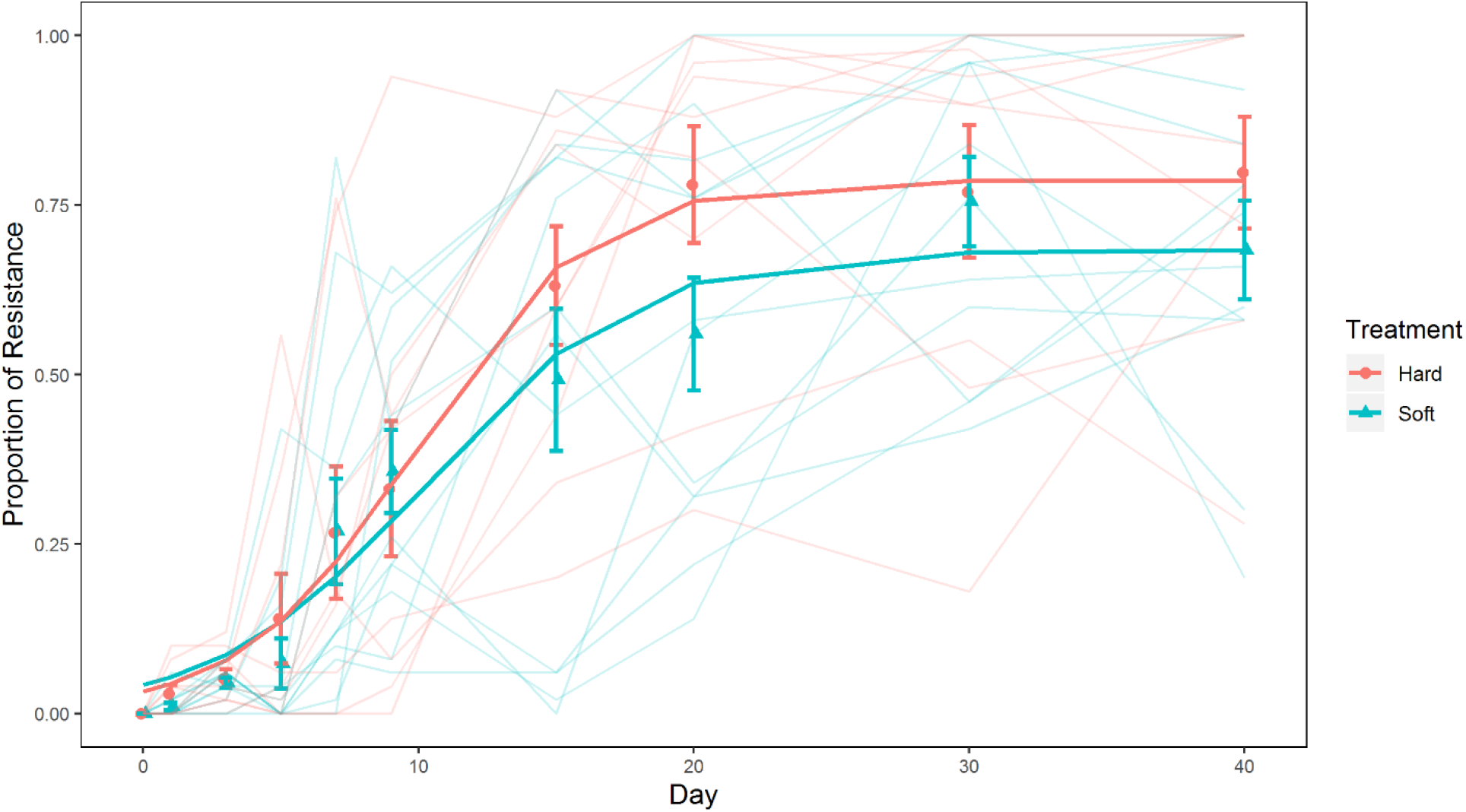
The proportion of resistant individuals. Each point is the mean proportion of resistant individuals from either soft selected (blue) or hard selected (red) population replicates (n=12). Error bars represent standard error. Lighter lines are the proportion of resistant isolates in each replicate population.

Prolonged coexistence between divergently adapted types in a spatially heterogeneous environment should be underlain by a fitness trade-off across conditions of growth. To evaluate this prediction, we surveyed the evolved populations that had evolved under divergent selection for fitness trade-offs between resistant and sensitive strains. We first measured the MIC and growth rate under drug-free conditions of four randomly chosen resistant and sensitive colonies, respectively, from each evolved population. We have previously shown that resistance, when it first evolves, is costly under drug free conditions (29) but this cost can be reduced by compensatory mutations that improve growth rate under permissive conditions without compromising resistance in a spatially variable environment (28). We see similar results here under both forms of population regulation: the initially strong trade-off between MIC and growth rate under permissive conditions is undetectable by day 40 due to the evolution of increased growth rate of resistant types in the absence of drug (**Fig 2**). Prolonged coexistence therefore cannot be due solely to costs associated with growth rate under permissive conditions. Second, we used competitive fitness assays against the ancestral strain under drug free conditions to provide a more direct estimate of the extent of adaptation for a pair of resistant and sensitive isolates from each evolved population (**Fig 3**). Resistant strains (filled circles) show a substantial fitness cost in the absence of drug at day 10 (HARD day 10 mean ± SE: 0.880 ± 0.037 [CI: 0.804, 0.957]; SOFT day 10 mean ± SE: 0.920 ± 0.037 [CI: 0.843, 0.996]), but not at day 40 under both forms of population regulation (HARD day 40 mean ± SE: 0.986 ± 0.041 [CI: 0.902, 1.07]; SOFT day 40 mean ± SE: 1.01 ± 0.038 [CI: 0.938, 1.09]). This result mirrors that seen in the growth rate assays and is consistent with the substitution of compensatory mutations that improve fitness under permissive conditions without compromising resistance. Sensitive strains also show improvements in fitness in the absence of drug, although the dynamics of this process differ for soft and hard selection. Under hard selection, sensitive strains initially adapt rapidly to drug-free conditions (HARD Day 10 mean ± SE: 1.19 ± 0.043 [CI: 1.11, 1.26]), and show little evidence of further improvement by Day 40 (HARD Day 40 mean ± SE: 1.13 ± 0.041 [CI: 1.04, 1.21]; contrast day 10 – 40: 0.059 ± 0.045, ***P*=0.895**; **Fig 3**, red triangles). Sensitive strains under soft selection, on the other hand, adapt more slowly initially, though they continue to increase in fitness at rates that parallel the fitness increases seen in their coexisting resistant strains (contrast SOFT Sen – Res fitness increases mean ± SE: 0.001 ± 0.02, ***P*=0.725**). We attribute this difference in rate of adaptation in sensitive types to a reduced beneficial mutation supply rate – the product of the genome-wide beneficial mutation rate, *U*_*b*_ and the effective population size, *N*_*e*_ – under soft selection arising from the fixed contribution of individuals from each patch. Under hard selection, each patch contributes individuals to the dispersal pool in proportion to that patch’s productivity, which early in the experiment is by design heavily weighted towards sensitive types. Selection should thus be far more effective at generating adaptation amongst sensitive types under hard selection than it is under soft selection early in the experiment. This effect notwithstanding, sensitive types are always fitter under drug free conditions than resistant types at both time points, consistent with the idea that fitness trade-offs underpin coexistence between these two classes of phenotypes over the course of the experiment.

**Figure 2.**
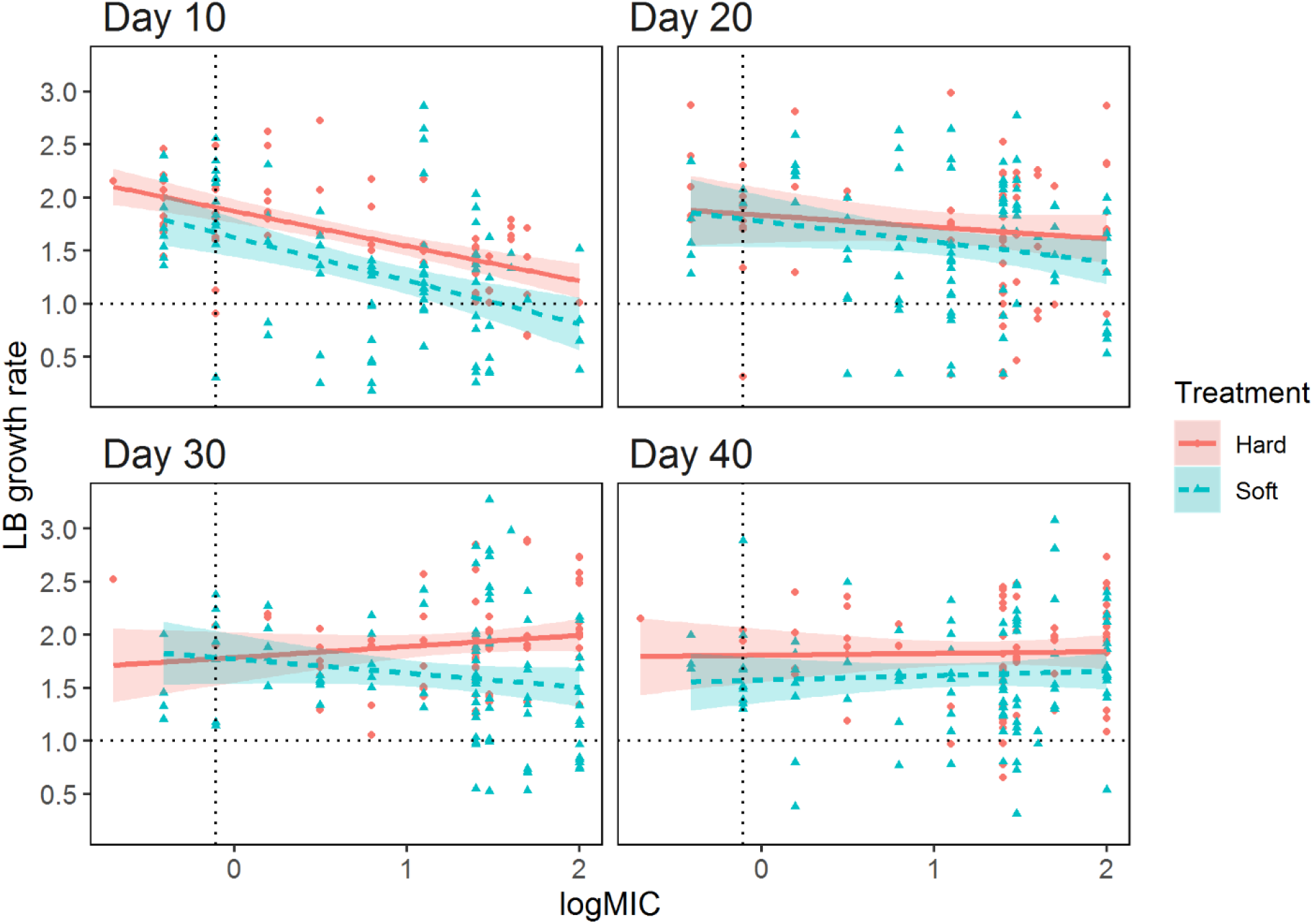
Trade-off between growth rate and log MIC. Each point represents an isolate taken from either a soft selected (n=96) or hard selected (n=72) population. Growth rates and log MIC are standardized to the ancestral strain, represented by the dashed horizontal and vertical lines, respectively. Shaded regions represent standard error.

**Figure 3.**
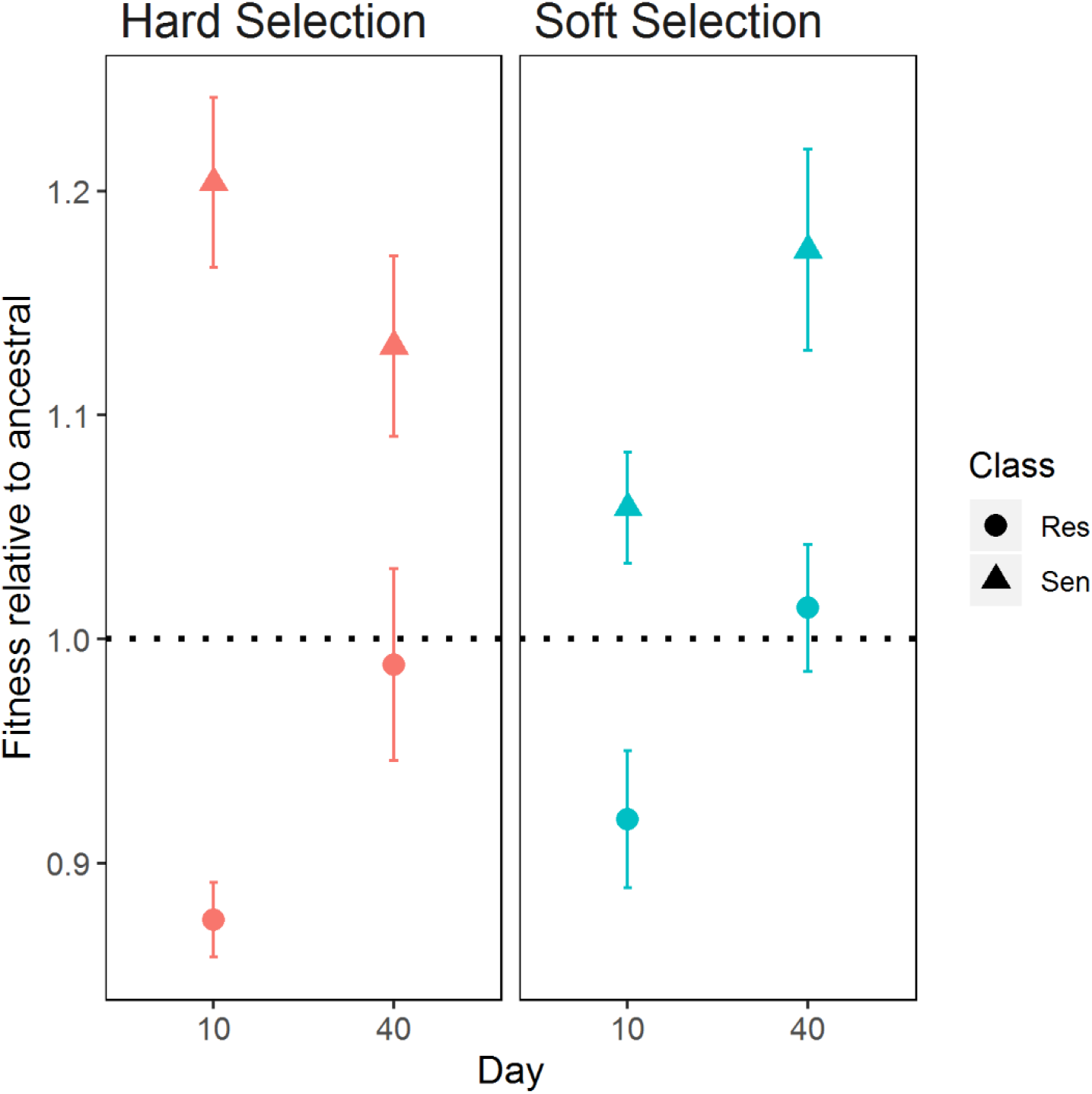
The relative fitness of resistant and sensitive isolates to ancestral. Each point represents mean relative fitness of resistant (circle) or sensitive (triangle) isolates relative to ancestral strains. Error bars represent standard error. Horizontal dotted line represents fitness of ancestral strains.

Is coexistence between resistant and sensitive types stable on evolutionary time scales? Theory suggests that diversity should be maintained by negative frequency-dependent selection under soft but not hard selection. To test this prediction, we assayed fitness in reciprocal invasion-from-rare competitions between pairs of resistant and sensitive isolates chosen at random from six populations evolving under soft or hard selection at day 10 and again at day 40 (6 pairs × 2 treatments × 2 time points = 24 sets of invasion-from-rare experiments). We find that, consistent with theory, fitness is negatively frequency-dependent under soft selection at both time points observed (**Fig 4**; SOFT day 10 slope ± SE: −0.199 ± 0.044 [CI: −0.287, −0.112]; SOFT day 40 – 0.203 ± 0.048 [CI: −0.300, −0.106]). Notably, the slope of the fitness functions at the two time points are statistically indistinguishable (***P*=0.958)**. This result suggests that diversity can be stably maintained over the long-term via soft selection.

**Figure 4.**
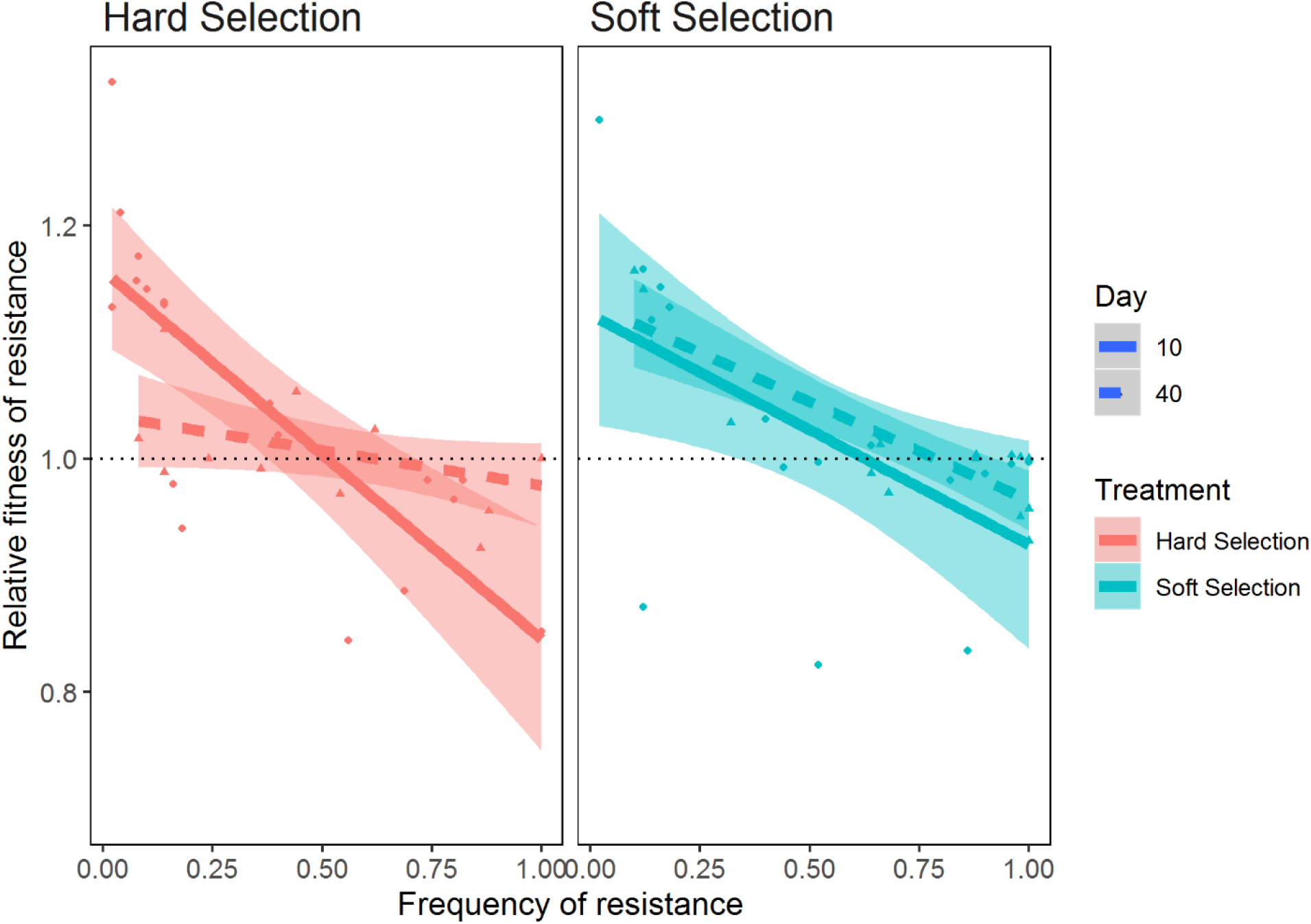
Negative frequency dependent selection. Each point represents the relative fitness of a resistant colony relative to a paired sensitive colony when plotted against its initial frequency day 10 (red) and day 40 (blue). Shaded regions represent standard errors. A signal for negative frequency dependent selection is given by the negative correlation between the relative fitness of resistant isolates and its initial frequency, and crosses the horizontal dotted line.

We also find evidence of negative frequency-dependent selection under hard selection at day 10 (−0.299 ± 0.046 [CI: −0392, −0.206]), a result that is contrary to predictions of theory but consistent with our previous work (28). We suspect negative frequency dependent selection is generated by the batch culture conditions used in this experiment: the evolution of resistance rapidly restores any initial imbalance in cell density between patches and allowing each sub-population to reach stationary phase before being transferred, resulting in local population regulation within each patch. By day 40, however, in line with theory, negative frequency dependent selection cannot be detected: the slope of the regression is both significantly different from day 10 (***P*=0.0016**) and indistinguishable from 0 (−0.071 ± 0.052 [CI: −0.175, 0.033]). Thus, the coexistence of resistant and sensitive strains under hard selection early in the experiment is only quasi-stable: the fitness function changes over time from being initially negative to effectively zero by the end of the experiment, and we expect diversity to be eventually lost from these populations as a result. That we see a significant enrichment of resistant strains under hard selection relative to soft selection by the end of the experiment is consistent with this prediction.

## Discussion

We have shown how two extreme forms of population regulation – at the level of the total, mixed population (hard selection) and at the level of spatially distinct patches (soft selection) – can impact the emergence and fate of diversity under strong divergent selection. Our leading results are that, while rates of diversification are not substantially affected by the scale at which population regulation occurs, the long-term fate of diversity is. Consistent with theory, diversity is more readily maintained when population regulation is local (soft selection) rather than global (hard selection). Moreover, the mechanism ensuring coexistence is negative frequency dependent selection, which while present initially in under both forms of population regulation, persists over the long-term only under soft selection. As a consequence, a polymorphism supported under soft selection can persist through time and is robust to continued evolution of each niche specialist type. By contrast, polymorphisms that initially emerge due to strong divergent selection under hard selection do not persist on evolutionary time scales, as they are eventually replaced by the emergence of a single, broadly adapted generalist type. Together, these results suggest that soft selection could be a general mechanism for the prolonged maintenance of ecological diversity on evolutionary time scales.

The inferential strength of this conclusion relies, of course, on the extent to which our experiment represents an accurate model of how divergent selection, dispersal, and population regulation operate in more natural settings. On this, we have very little empirical information, as we know little about the spatial scale at which divergent selection operates relative to dispersal in natural populations (30–32) and even less about how population size is regulated (33). Nevertheless, there could be situations where conditions similar to ours might be met. One possibility is selection imposed by the use of distinct antibiotics on different wards of a hospital (28). Selection for resistance, in this case, would be divergent across wards and dispersal would occur through the movement of microbes by patients, hospital staff, and others among wards. How population size is regulated in this example remains an open question, and one that has important consequences for efforts to manage or control the evolution and spread of resistance under different dosing regimes. If populations are regulated at the level of the ward, in a manner resembling soft selection, then using distinct drugs on different wards could preserve the effectiveness of antibiotic therapy by preventing a single, multi-drug resistant type from evolving. If, on the other hand, population regulation is more like hard selection, with some wards being more productive than others in terms of the total number of resistant bacteria dispersing among wards, a multi-drug resistant is expected to evolve eventually, although it could take longer to evolve than under constant or temporal selection (Leale and Kassen 2018).

Negative frequency dependent selection represents a powerful mechanism to maintain variation within populations, as demonstrated by experiments in controlled settings (34) and in natural populations(35,36). That said, our results suggest that it would be inappropriate to see the stable maintenance of diversity through negative frequency-dependent selection as somehow freezing or halting the evolutionary process altogether. The strong divergent selection supporting ecological differentiation between resistant and sensitive types in our experiment does not guarantee their prolonged coexistence under hard selection, nor does it prevent continued adaptation of each niche specialist type under soft selection. Indeed, the continued adaptation of niche specialists in the presence of extensive diversity has been reported in other systems (37,38). These results suggest that the conditions for coexistence can themselves evolve and that models for the maintenance of ecological diversity on evolutionary time scales need to account for such continued evolution. More generally, the conditions promoting diversification may be quite distinct form those governing its long-term maintenance. To the extent that the model communities that we have studied in the lab capture the essential features of how selection works in nature, our results point to a world that is far more genetically and ecologically dynamic than traditional views, which have often relied on equilibrium analyses to interpret diversity, would have it.

## Authors’ contributions

Experimental design, data analysis and writing was completed by R.K. and P.J.C. Experiments were performed by P.J.C.

## Acknowledgements

We thank Alanna Leale for providing insight regarding experiment design, as well as the summer students Eric Roberts, Mohammed Arab and Kayla Garvey for their help in the laboratory.

## Methods

### Experimental evolution

A single colony of *P. aeruginosa* strain PA14, and an isogenic strain containing a *lacZ* insertion was grown overnight in lysogeny broth (LB: bacto-tryptone 10g/L, yeast extract 5g/L, NaCL 10g/L) and used to found a total of 72 replicate populations by adding 15uL of culture into 1.5ml of fresh media. Populations were propagated every 24 hours by adding a 15 uL aliquot from a mixed culture into 1500 uL of fresh media in 24-well microtiter plates. Populations were incubated and agitated using an orbital shaker (150 rpm) at 37°C for 40 days, or ~264 bacterial generations. Colonies possessing the *lacZ* insertion appear blue when cultured on agar plates supplemented with 40 mg L^−1^ of 5-bromo-4-chloro-3-indolyl-beta-D-galactopyranoside (X-Gal), and are visually distinct from the wild type white colouration. Populations were routinely screened every 3-4 days (20-26 generations) for cross contamination before being archived in a 16% glycerol solution at −80°C.

The experiment was designed such that 24 replicate populations would be distributed among homogeneous environments of permissive LB media, homogeneous environments of LB supplemented with 0.3 μg/ml of the fluoroquinolone antibiotic ciprofloxacin, and heterogeneous environments where one subpopulation was cultured in LB and the other in LB supplemented with ciprofloxacin. This concentration of ciprofloxacin was previously determined to reduce the maximal growth of a sensitive PA14 ancestor to 20% of full growth in LB over a 24h period. Within each environment, 12 replicate populations would undergo either soft or hard selective regimes. Soft selection was imposed by regulating the cell densities of subpopulations prior to population mixing and redistribution into each patch. This was done by using optical density (OD), measured at 600nm using a spectrophotometer (Elx-800: BioTek Instruments Inc., Winooski, VT), as a proxy for cell density. We regulated cell densities from each patch by diluting the denser sub-population with unsupplemented LB until the OD was similar to the complementary patch. Hard selection was imposed by mixing a fixed volume (500 μl) aliquot from each subpopulation before redistribution. All mixed populations from both selection regimes were diluted to a common density of approximately 0.45 OD prior to transfer to fresh media.

### The evolution and spread of resistance

We assayed the frequency of resistance in the mixed samples from both sub-populations of a given lineage (12 lines from soft selection, 9 lines from hard selection) on days 1, 3, 5, 10, 15, 20, 30 and 40 by isolating 50 random colonies from LB plates and then streaking these on agar plates containing 2μg/ml ciprofloxacin. Plates were visually inspected after 24 hours growth at 37°C.

### Trade-offs and costs of resistance

We assayed for fitness trade-offs for antibiotic resistance by measuring the minimum inhibitory concentration (MIC) and the maximal growth rate of eight isolates, of which four were randomly selected resistant and four were randomly selected sensitive colonies, from each evolved population (12 lines from soft selection, 9 from hard selection). MIC was determined by incubating 100ul of overnight culture in 100ul of LB supplemented with concentrations of 0, 0.2, 0.4, 0.8, 1.6, 3, 6, 12, 25, 30, 40, 50ug/ml ciprofloxacin respectively, and observing for growth after 24 hours. Growth rate of the same eight isolates was estimated by inoculating triplicates of 5ul of overnight culture into 195 of LB and measuring the OD at 90 minute intervals over 24 hours. Gen5 software (BioTek Instruments Inc., Winooski, VT) was used to calculate the maximal growth rate.

In addition, we further examined the costs of resistance by estimating the fitness of a randomly selected sensitive and resistant isolate from each line (6 lines from each soft selection and hard selection) on day 10 and day 40 in competition against the PA14 common ancestor in LB medium. Overnight LB-grown cultures of each evolved isolate and the reciprocally marked ancestor were mixed 1:1 by volume and then inoculated into LB medium. We tracked the change in relative abundance of the evolved isolate and ancestral strain over 24 hours, plating the mixed culture at 0 hrs and 24hrs and counting the relative frequency of each type by visual inspection on X-gal-supplemented plates. The relative fitness (***ω***) of evolved isolates was estimated using the equation:

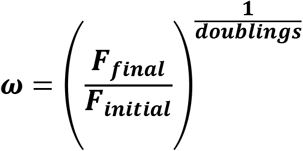

where F_initial_ and F_final_ represent the ratios of the frequency of the evolved population to the frequency of the marked ancestor at 0 and 24 hrs, respectively. Doublings refers to the number of generations between the initial and final measurements (approximately 6.6 generations).

### Negative frequency dependent selection

We estimated the strength of negative frequency-dependent selection in each lineage using a reciprocal competitive invasion experiment between randomly chosen resistant and sensitive isolates at days 10 and day 40. Overnight cultures of each isolate were mixed at volumetric ratios of 1:9, 1:1 and 9:1, following the protocol of Leale and Kassen (2018) and allowed to compete for two transfer cycles (48hrs) under the same population regulation protocol from which the isolates were taken. We estimated the frequency of resistant colonies by plating at least 50 random individual colonies prior to, and after, the competition on LB agar plates supplemented with ciprofloxacin at a concentration of 2μg/ml. The relative fitness of resistant isolates was estimated using the equation:

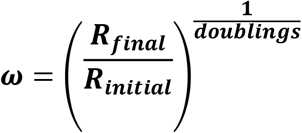

where R_initial_ and R_final_ represent the ratios of the frequency of resistant types to the frequency of sensitive types, before and after the competition, respectively. Doubling refers to the number of generations between the initial and final measurements (approximately 13 generations; ~6.6 generations per 24h cycle).

### Statistical Analysis

All statistical analyses were conducted using R statistical software (39). We modelled the evolution of resistance using a two parameter logistic growth model using nonlinear least squares (NLS) (40). Using an analogous comparison between the evolution of resistance and a logistic growth model, we generated an estimate of the maximal rate at which resistance propagated throughout each individual population and the maximum proportion of resistance individuals at the end of the experiment. Comparisons of the final proportion of resistance between the selection treatments were done using a generalized linear mixed model (GLMER) (41) with a binomial link, with a random intercept of individual population to account for resampling across time (repeated measures). Contrasts of maximal growth rates were compared using a linear model, with treatment being used as a categorical factor.

We measured the strength of negative frequency dependent selection at two timepoints using reciprocal invasion experiments of a resistant-sensitive pair taken from each independent population. We calculated and modelled the relative fitness (*ω*) of the resistant individual as a function of the initial frequency of resistance. Evidence for negative frequency dependence would be given if there is a statistically significant negative correlation between the relative fitness of resistant isolates and the initial frequency of resistance.

We also tested for the presence and strength of trade-offs within each selection regime by calculating the relative fitness of resistant or sensitive isolates when competed against an ancestral strain. The relative fitness of a resistant or sensitive individual was modeled against the categorical factor of “treatment” with a random intercept of individual population to account for repeated measures.

