## Supplementary figures and images for "The Evolution and Fate of Diversity Under Hard and Soft Selection"

### Supplementary Fig 1.tif

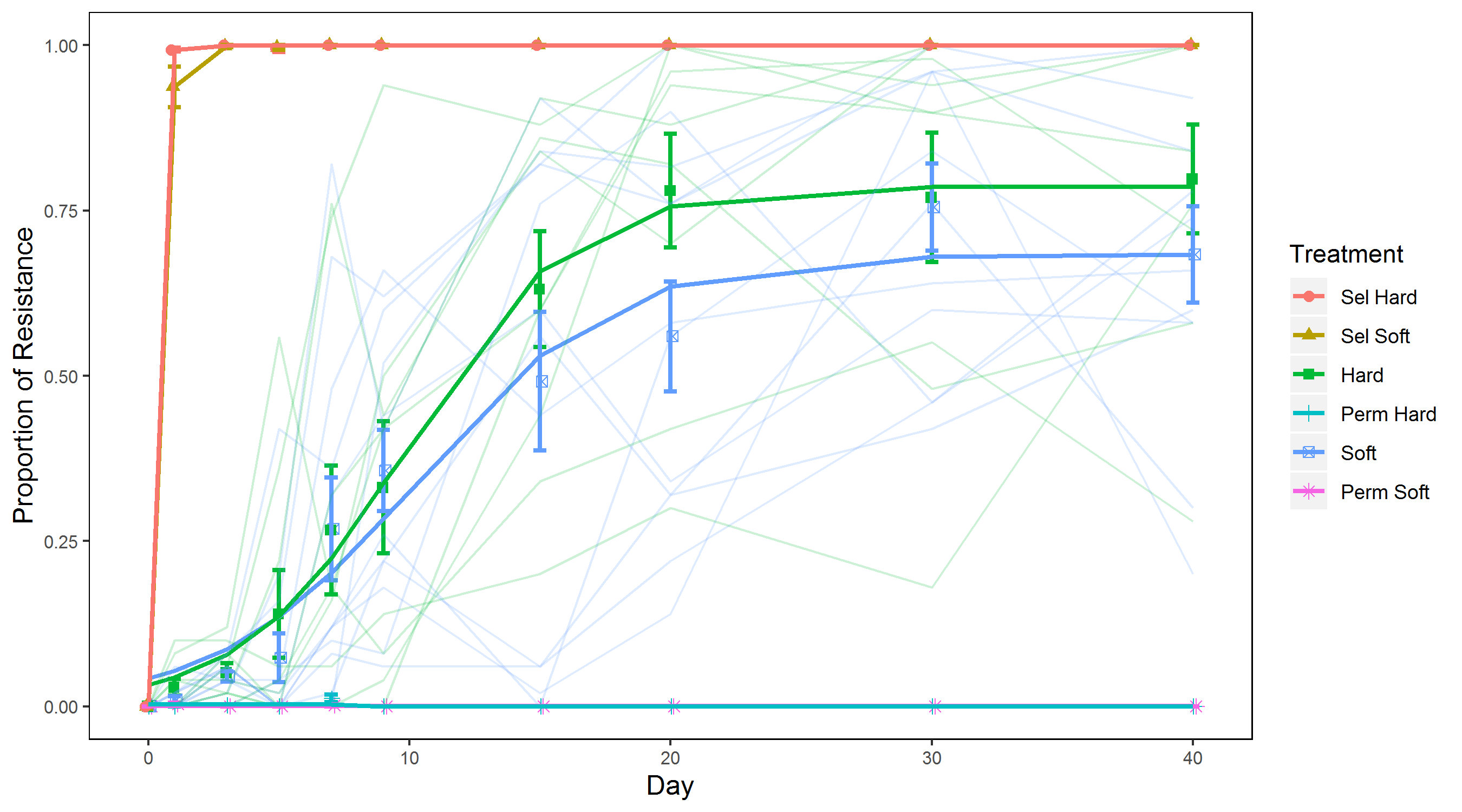
